# Contrasting mechanisms for using humidity as cue for seasonal polyphenism in two tropical butterflies

**DOI:** 10.1101/2025.10.14.682489

**Authors:** Tarunkishwor Yumnam, Freerk Molleman, Urszula Walczak, Ullasa Kodandaramaiah

## Abstract

Many tropical butterflies show wing pattern plasticity across dry and wet seasons. Dry-season morphs of satyrines (Nymphalidae: Satyrinae) typically are cryptic with small or no eyespots on the ventral wing surface. In contrast, wet-season morphs have large eyespots on their wing margins. These eyespots deflect predator attacks away from vital body parts. Extensive studies have shown that eyespot size is influenced by the temperature experienced during larval development. Temperature-mediated eyespot size plasticity is adaptive in regions where temperature increases with the onset of the wet season. However, many tropical regions lack a reliable temperature-rainfall correlation, and thus, temperature may not always be a reliable predictor of the upcoming adult environment. A potential environmental cue that correlates strongly with seasonality is relative humidity (hereafter, humidity). With the approach of the wet season, humidity increases, and then declines with the onset of the dry season. Here, we tested whether humidity modulates eyespot-size plasticity in two sympatric butterflies that experience similar ecological pressures: *Mycalesis mineus* and *Melanitis leda*. We found that humidity directly influenced eyespot size in *My. mineus*; individuals reared at high humidity developed into adults with larger eyespots than those from low humidity. In contrast, eyespot size in *Me. leda* showed a weak direct effect of humidity. Instead, humidity modulated eyespot size in *Me. leda* indirectly through induced changes in the host plant quality. Since the late larval and prepupal stages are temporally closer to the adult environment, selection may favour greater sensitivity to the environmental cue during these stages than during early larval stages. To determine the humidity-sensitive phase, we conducted a switching experiment where humidity was shifted from low to high, and *vice versa*, at different developmental stages. We demonstrated that, in *My. mineus*, wandering larval and prepupal stages are sensitive to humidity, whereas the early-feeding larval phases are insensitive. Contrastingly, humidity switching did not influence the eyespot size in *Me. leda*. This study highlights how two diverging species, evolving under similar ecological pressures, integrate environmental cues in distinct ways to modulate wing pattern plasticity.

## Introduction

Many organisms in seasonal environments use phenotypic plasticity to produce phenotypes that fit the season (1), but this may make them vulnerable if cues become unreliable under climate change. In short-lived organisms, such seasonal polyphenism is often achieved through developmental plasticity in which cues experienced during immature stages are used to develop fitting adult phenotypes (2). These cues need to be both reliable for predicting the environment of the adult, and perceivable by the immature organisms. Which cues are reliable and perceivable will depend on the region, with, for example, photoperiod being favoured in more temperate regions where day length changes markedly with the seasons (e.g. 3), while tropical species may need to rely on cues such as temperature and humidity (e.g. 4) to predict wet and dry seasons. Even though seasonal polyphenism is often observed in the wild, we have a limited understanding of which cues are used and how, and data for too few species to trace their evolution (but see 5,6).

A prominent example of adaptive seasonal plasticity is the widespread polyphenism of wing patterning exhibited by many tropical satyrine (Nymphalidae; Satyrinae) butterflies (e.g., 5,7– 13). Adults of these butterflies that develop during the wet season have large, conspicuous ventral eyespots on the wing margin, while the dry-season adults have significantly reduced, or no eyespots, with overall cryptic ventral colouration. The large eyespots of the wet-season morph can deflect predator attacks away from vital body parts and misdirect towards the less vital wing marginal areas (14–17). In contrast, the dry-season wing patterns help to camouflage the butterflies against the dry vegetation (17,18).

A large number of studies have suggested that temperature during developing stages regulates eyespot-size plasticity in these tropical satyrines (7–9,19,20). Temperature-mediated eyespot plasticity has been experimentally confirmed in several African and Asian satyrines (4,21–27). This temperature-regulated plasticity has been especially well-characterized in the African butterfly *Bicyclus anynana*, where lower temperature during development results in dry-season adults with small eyespots, and high temperature leads to wet-season adults with large eyespots (13,19,21,22,28–30). Eyespot expression is mediated by ecdysteroid hormonal regulation induced by the temperature during development (22,30,31). The temperature-sensitive period of *B. anynana* occurs from the late larval stage to the early pupal stage (29,30), after larval feeding has ceased, but wings are not fully formed yet. Therefore, research so far overwhelmingly supports the idea that eyespot plasticity is regulated by temperature.

However, temperature cannot be the only cue used by these butterflies. The use of temperature as the cue for the regulation of eyespot size is expected to be adaptive if temperature experienced during the immature stages reliably predicts the season in which adults fly. Studies on the developmental control of eyespot size in *B. anynana* have been conducted almost exclusively on a single population sourced from Malawi. In Malawi, the dry seasons are cooler than the wet seasons, and therefore, temperature-regulated eyespot plasticity is likely to be adaptive (8,29,32). However, many species, including *B. safitza* (27,33), *Mycalesis mineus* (L. 1758) and *My. perseoides* (26) exhibit eyespot size plasticity, but also occur in regions where there is no clear temporal correlation between temperature and rainfall. Thus, in many populations with eyespot size plasticity, the temperature experienced by larvae may not reliably predict the season in which adults live, and selection would thus not favor using temperature as a cue. Indeed, temperature does not influence eyespot size in populations of multiple *Bicyclus* species in south-western Africa (34), where the dry season is warmer than the wet season. Therefore, temperature-regulated eyespot plasticity is unlikely to be adaptive in many butterfly species or populations, and these butterflies are expected to rely on other cues (11,34,35).

Two variables that differ strongly between dry and wet seasons in the tropics are precipitation and humidity. Precipitation and humidity are positively correlated in most parts of the tropics (36,37), including tropical continents (38) and islands (39). For instance, in our study location, in southern India, the wet season begins in May and ends in November, while the dry season spans December to April (Fig. 1). Although similar temperature ranges occur during both seasons, the absolute humidity and relative humidity (hereafter, referred to as humidity) are, respectively, over 20 g/m^3^ and 80% during large part of the wet season, reaching maximum of 22.3 g/m^3^ and 89.5% during wet season and minimum of 16.9 g/m^3^ and 63.5% during the dry season (Fig. 1). Thus, humidity experienced by larvae may reliably predict whether the emerging adults will face the dry-or the wet-season environment. Humidity has been shown to affect pupal coloration in *My. mineus* (one of our study species) (40), suggesting that immatures are able to respond to humidity. However, humidity has not been experimentally demonstrated to influence eyespot size in any butterfly (but see suggestion for an effect of humidity in *Melanitis leda* (L. 1758) in (4)). Here, we test whether humidity experienced during development can regulate adult eyespot size in two distantly related satyrine butterflies.

**Figure 1.**
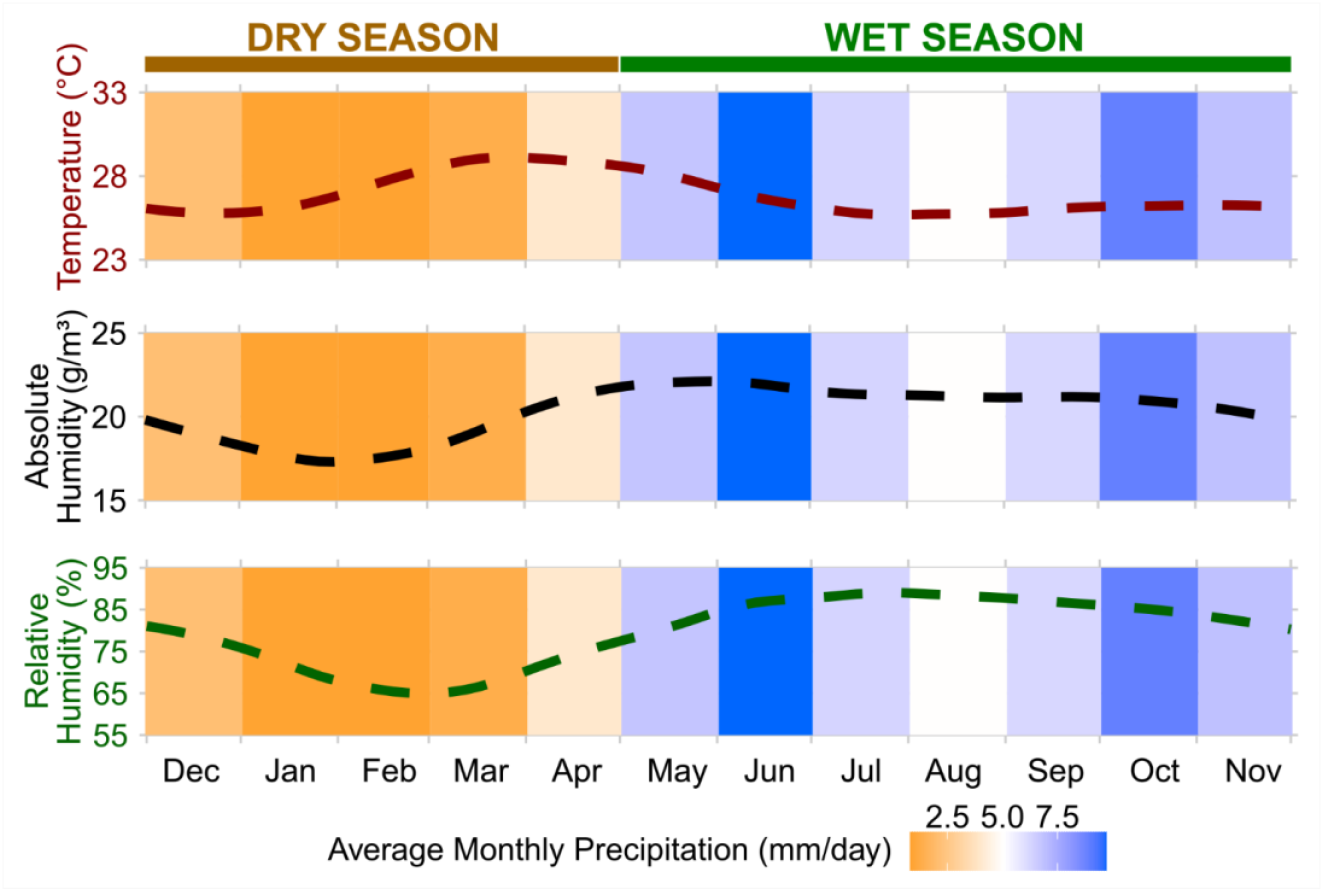
Smoothed curves of daily averages of relative humidity (dark green), absolute humidity (black), temperature (dark red) at 2 meters above the ground, and monthly average of precipitation (orange-blue gradient bars) for the years 1981-2024 at the study location, Vithura, India (8°40’56.07” N, 77°8’8.36” E). Data were sourced from NASA MERRA2 (55).

Larvae may directly sense the level of humidity, or the effect may be via the host plant. Although it is unclear whether insects can sense precipitation directly, larvae can probably sense humidity levels through hygrosensory receptor neurons located in sensory hairs or sensilla (reviewed in (41)). Thus, larvae or prepupae of butterflies may be able to sense the differences in humidity between seasons and translate this information to developmentally regulate eyespot size. Butterflies may also detect variation in host plant traits that are induced by humidity. Satyrine butterfly larvae typically feed on multiple grass species (23,42,43). They may feed on different species during different seasons because the community of grasses available can differ between seasons (44). Host plant species have been shown to influence eyespot size (4,23,45). It seems likely that host plant species affect eyespot size via differences in nutritional quality, as there is a correlation between larval development time and adult phenotype across plant species (4,23). In natural habitats, within-species host plant quality of grass generally declines as the wet season transitions to the dry season (46,47). Thus, the quality of the same host plant may differ between seasons and can also affect the eyespot size. Kooi et al. (48) showed that *B. anynana* larvae fed on drought-stressed plants develop into adults with smaller eyespots compared to adults developing from larvae fed on well-watered plants. However, these studies on how host plant affects eyespot size did not provide data on host plant quality. Therefore, we tested whether humidity can affect host plant quality, and if this has a cascading effect on butterfly eyespot size.

To test if and how humidity affects eyespot size, we performed experiments on sympatric populations of two tropical satyrines *My. mineus* and *Me. leda*. These species represent lineages that diverged more than 50 million years ago (49), but face similar ecological pressures today. Larvae of both species utilize grasses as host plants (23,42,43). Firstly, we reared larval hatchlings at two humidity conditions representing the dry and wet seasons in their native habitats. Secondly, we performed switching experiments in which larvae were switched between levels of humidity at two different developmental stages (wandering and prepupal stages) to test if the response to humidity occurs during or after the larval feeding stages. Finally, to test whether butterfly eyespot size is sensitive to humidity-mediated differences in host plant quality, we also reared larvae on plants that were maintained at either low or high humidity, while larvae themselves were kept under low humidity. To quantify the host plant quality, we measured the carbon to nitrogen (C:N) ratio. High C:N in the host plants is known to adversely affect herbivorous insect development (reviewed in (50)). Moreover, Nitrogen is assimilated into amino acids, and increased amino acid availability can activate the insulin/TOR signalling pathway (51,52), which subsequently leads to the production of 20-hydroxyecdysone or 20E (reviewed in (53)) – the regulator of eyespot size in *B. anynana* (5).

If butterflies use humidity experienced during their larval stages as a cue to modulate eyespot size, individuals reared at low humidity should develop into adults with smaller eyespots than those reared at high humidity. Since eyespot development is known to start at the early pupal stage (54), any environmental cue experienced after the onset of pupation is unlikely to affect the eyespot development. Compared to the early larval stage, the late larval stage and prepupal stages are temporally closer to the upcoming environment that the eclosing adults will experience. Thus, selection is expected to favour sensitivity to environmental cues during the latter two stages. Therefore, we predicted that the larval wandering stage, i.e., the stage immediately prior to pupation, and the prepupal stage are sensitive, whereas the early larval stage is insensitive. Alternatively, larvae may not respond directly to differences in ambient humidity, but sense differences between plants exposed to low and high moisture, and use these host-plant cues to regulate eyespot size. If so, individuals developing on well-watered plants should have larger eyespots compared to those developing on drought-stressed plants when reared under the same conditions.

## Methods

### General procedure and data collection

Wild adults (ca. 35 females each) of *My. mineus* and *Me. leda* were caught from within a 5-km radius of the Indian Institute of Science Education and Research Thiruvananthapuram (8°40’56.07” N, 77°8’8.36” E) campus using banana-baited traps, and housed in aluminium cages (45 cm x 40 cm x 40cm). Adults were provided ripe banana slices as food, and ca. two-weeks-old maize (*Zea mays*, dent corn variety) for oviposition. The maize host plants were grown in plastic pots (9 cm x 10cm x 10cm) in a semi-natural setting inside an outdoor enclosure fitted with sprinklers for watering at regular intervals. Hatchling larvae were collected daily, and randomly assigned to experimental treatments within 24 hours of hatching. The larvae were reared inside nylon mesh sleeves (0.135 m x 0.28 m x 0.95 m) at a density of 20 (*My. mineus*) or 15 (*Me. leda*) per sleeve. All larvae within a sleeve hatched on the same day. A single pot with two-weeks-old (± two days) maize plants was placed as larval food in the sleeves. The plants were replaced at regular intervals and watered daily (hereafter, referred to as well-watered plants), except in the case of the *60lowhost* treatment, which included drought-stressed plants (described in more detail in subsection *Treatments*). For all sleeves, larval host plants were replaced every third day until the sixth day, and every alternate day thereafter. The sleeves with larvae and host plants were placed inside two growth chambers (model: E-36VL, Percival Scientific, Iowa, USA) set at 25°C, 12:12 hours light:dark photoperiod, and either 60% or 85% humidity. The two humidity values were chosen to reflect the natural variation of humidity during dry and wet seasons in the location where the butterflies were sourced from (Fig. 1). To eliminate potential growth-chamber effects, the humidity setting of the growth chambers was alternated periodically.

Once fifth instar larvae were detected in a sleeve, the sleeve was inspected daily for the presence of wandering-stage larvae. Such larvae were collected and placed in 100 ml transparent containers (Tarsons, Kolkata, India) covered with black fine-mesh on the top, until eclosion (more details in subsection *Effects of humidity and Determination of humidity-sensitive stage)*. The containers were checked daily, and the dates of pupation and eclosion were recorded. Pupal mass was measured 24-48 hours post-pupation. For *My. mineus*, sexing was done during the adult stage as males are easily distinguishable from females by the presence of androconial brands on the wing (56). *Me. leda* were sexed during the pupal stage following Prasannakumar et al. (23). Adults were euthanised by freezing and their wings excised. Wings were photographed inside a light box and against a grey standard with scale, using a Nikon D3200 (Nikon, Tokyo, Japan) digital camera fitted with a 105 mm fixed focal length macro lens (Sigma Corporation, Kanagawa, Japan). Wing images were used to measure wing and eyespot size.

### Treatments

The experiments for both species included seven treatments that were run in parallel. The treatments were designed with *a priori* planned comparisons to test whether humidity, humidity switching during different life stages, and humidity-induced host plant quality influence eyespot size.

1. ***60%***: Reared continuously at 60% humidity from hatching until adult eclosion.
2. ***85%***: Reared continuously at 85% humidity from hatching until adult eclosion.
3. ***60to85ws***: Reared at 60% humidity from hatching and switched to 85% humidity from the onset of the larval wandering stage.
4. ***60to85pp***: Reared at 60% humidity from hatching and switched to 85% humidity from the onset of the prepupal stage.
5. ***85to60ws***: Reared at 85% humidity from hatching and switched to 60% humidity from the onset of the larval wandering stage.
6. ***85to60pp***: Reared at 85% humidity from hatching and switched to 60% humidity from the onset of the prepupal stage.
7. ***60lowhost***: Fed on drought-stressed plants and reared at 60% humidity until adult eclosion. Drought-stressed plants were those that were watered daily for two weeks, and then placed in a growth chamber at 60% humidity without watering for three days immediately before being given to larvae in the experimental sleeves.

### Quantification of plant quality

In order to test how moisture availability affected host plant quality, we quantified the carbon to nitrogen (C:N) ratio with a CHNS analyser, vario MACRO cube (Elementar, Langenselbold, Germany). C:N ratio was analyzed using uneaten leaves of plants that were fed on by larvae in sleeves for two days. Fifteen to twenty 20 leaves per pot from 9 randomly selected pots of host plant from ‘*60lowhost*’ and 8 pots each from ‘*60%*’ and ‘*85%*’ treatments were collected. The samples were dried for seven days at 27°C and then oven-dried at 60°C for 48 hours (57). The dried samples were ground and ca. 2 mg of the powdered samples were loaded in the CNHS analyser to estimate the percentage weight of C and N.

### Wing size and eyespot size measurement

Eyespot, wing shape and wing size were measured using a custom-written macro in ImageJ *v. 1*.*54g* (58). For both species, the forewing Cu1 eyespot size was quantified by measuring the diameters of the outer black ring and assuming that the eyespot was circular (4,59). Since the Cu1 hindwing eyespot of the wet-season morph is elliptical rather than circular, its size was quantified by assuming an ellipse using the longest (roughly parallel to the Cu1 vein) and shortest (roughly perpendicular to the vein and passing through the pupil) diameters. The distance between the end of the M1 vein and the meeting point of the Cu1 vein and the cell on the forewing was measured as a proxy for wingspan. Triangular areas on the forewing and hindwing were measured using fixed landmarks, as described in Prasannakumar et al. (23). These areas were used as proxies for wing sizes. For all analyses, eyespot size relative to the wing size (eyespot size divided by wing size) was used to account for size variation across individuals (4,60).

### Statistical analyses

All statistical analyses were performed using R *v. 4*.*4*.*2* (61) through RStudio *v. 2024*.*9*.*1*.*394* (62), and all the plots were generated using the R package *ggplot2 v. 3*.*5*.*1* (63). Analyses were done independently for each species and for the two wing surfaces (fore- and hindwings). For each planned comparison, generalized linear model (GLM) fitting was implemented using the R package *glmmTMB v. 1*.*1*.*11* (64) with relative eyespot size as the response variable, and treatment and sex as the explanatory variables. Relative eyespot size ranged continuously between 0.00185 and 0.4771 in *My. mineus*, and between 0.00212 and 0.1756 in *Me. leda*. Thus, models were fitted by applying a beta distribution with a logit link function. The global model included two-way interactions of all explanatory variables. Model elimination was performed using the *dredge* function in the package *MuMIn v. 1*.*48*.*4* (65), and the best-fitting model was chosen based on the lowest AICc value. The best-fitting model was also confirmed through likelihood ratio tests via the *anova* function. The generalized linear model analyses were followed by Tukey tests for *post hoc* analysis using the package *emmeans v*.*1*.*10*.*5* (66). The following comparisons were made.

### Comparison 1: Effect of humidity on eyespot size

To determine the overall effect of humidity on eyespot size in the two species, the treatments where larvae were grown at either low (*60%*) or high (*85%*) humidity from larval hatching to adult eclosion were compared. The effect of humidity could be direct cue use or via the plant, as plants were also exposed to the humidity treatment while being fed to larvae (plants were replaced every two days during most of the larval period).

### Comparisons 2a and 2b: Determination of humidity-sensitive stage

Two comparisons were done to determine if the humidity-sensitive stage occurs during or after larval feeding. Comparison 2a compared eyespot size across treatments in which larvae were all initially reared at low humidity, but either remained at low humidity (*60%)*, or switched to high humidity at the wandering (*60to85ws*), or prepupal stages (*60to85pp*) till adult eclosion. Similarly, Comparison 2b compared treatments *85%, 85to60pp* and *85to60ws*, where larvae were initially reared at high humidity, but either continued at this humidity or were switched to low humidity as wandering larvae or as prepupae.

### Comparison 3: Effect of humidity via host plant

This comparison was done to test whether humidity indirectly affects eyespot size via induced changes in the host plant. Eyespot sizes were compared across three treatments - *60lowhost, 60%* and *85%*. The C:N ratio of the host plants used in *60lowhost, 60%*, and *85%* was compared using a Kruskal-Wallis test implemented using the *kruskal*.*test* function. *Post hoc* comparisons between the treatments were performed using Dunn’s test with Holm correction using the *dunnTest* function in the R package *FSA v. 0*.*10*.*0* (67).

## Results

### Comparison 1: Effect of humidity

#### i) Mycalesis mineus

For the relative forewing as well as the hindwing eyespot size of *My. mineus*, the best-fitting GLM included treatment as the sole explanatory variable. Eyespot size was greater at *85%* than at *60%* humidity (Forewing: estimate = 0.711, standard error = 0.130, p < 0.0001, Fig. 2A; Hindwing: estimate = 0.626, standard error = 0.089, p < 0.0001, Fig. 2B).

**Figure 2.**
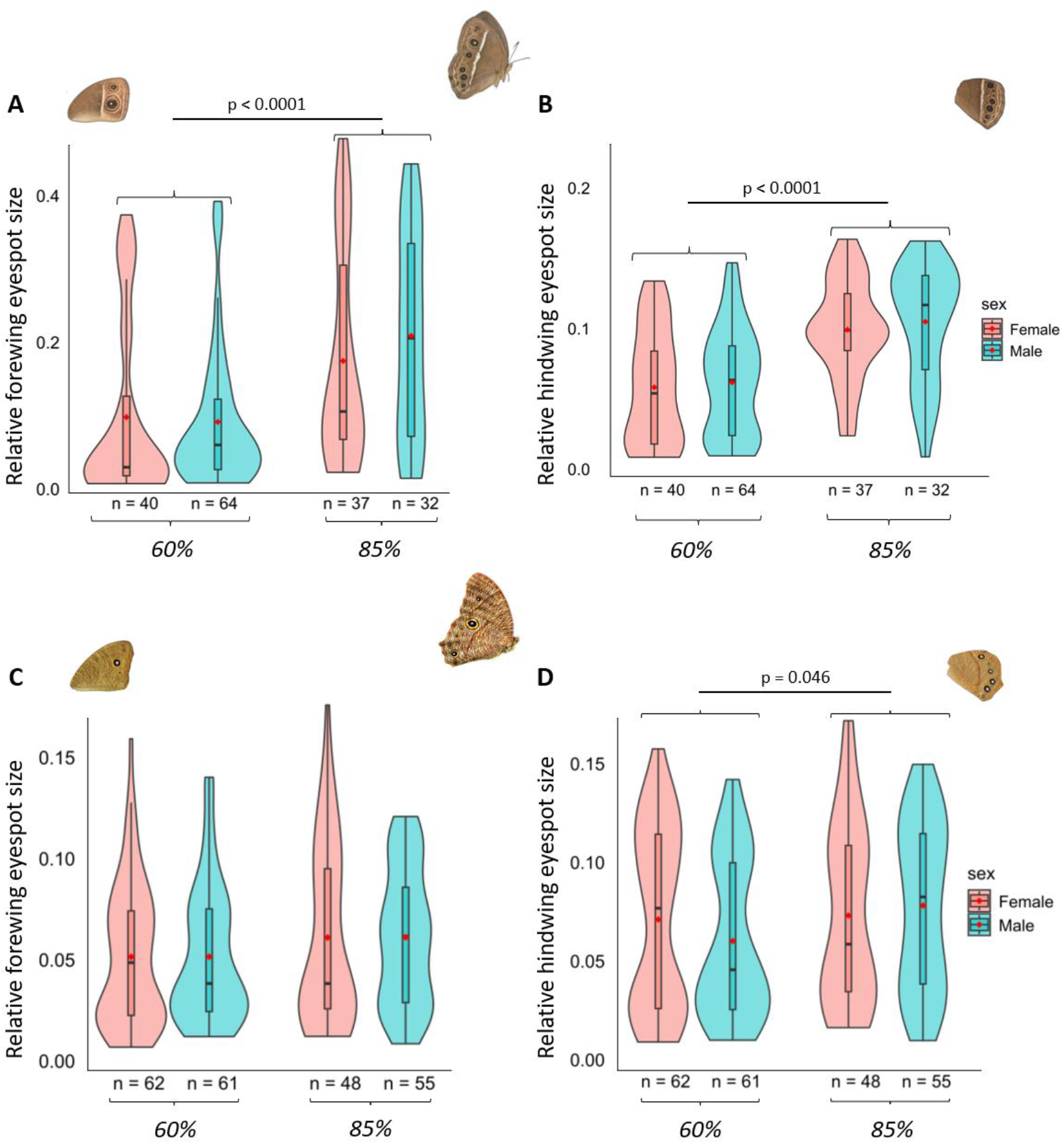
Violin plots showing the effects of humidity on the relative eyespot sizes of (A) forewing and (B) hindwing in *My. mineus*, and (C) forewing and (D) hindwing in *Me. leda*. The boxes inside the violin represent the 75^th^ (upper horizontal lines) and 25^th^ (lower horizontal lines) percentiles of the samples and medians (middle horizontal lines). The whisker lines indicate 1.5 times the interquartile range below and above the 25^th^ and 75^th^ percentiles, respectively. The red circles represent the respective averages. Shaded regions surrounding the boxes depict rotated kernel density plots. P-values reported are based on Tukey’s *post hoc* tests. Only significant differences between treatments are shown.

#### ii) Melanitis leda

The null model was the best-fitting model for the forewing in *Me. leda*. The relative forewing eyespot size was not significantly affected by humidity (Fig. 2C; coefficients from the next best model (ΔAICc = 1.047): estimate = 0.145, standard error = 0.082, p = 0.077). However, the best-fitting model for hindwing eyespot size contained treatment as the sole explanatory variable. Hindwing eyespot size was larger at *85%* than at *60%* humidity (Fig. 2D; estimate = 0.168, standard error = 0.084, p = 0.046).

### Comparisons 2a and 2b: Determination of humidity-sensitive stage

#### i) Mycalesis mineus

For forewing eyespots, the best-fitting model to test the effect of switching humidity from 60% to 85% (Comparison 2a) had treatment as the only explanatory variable (Table S1). Forewing eyespot size was larger when larvae were switched from 60% to 85% humidity at the prepupal stage (*60to85pp*) than those that remained in the 60% humidity treatment (Fig. 3A; Tukey’s *post hoc* test: p = 0.029). However, there was no effect of humidity switching at the wandering stage on forewing eyespot size (Fig. 3A; p-value based on Tukey’s *post hoc* tests: *60%* vs *60to85ws* = 0.488). In the case of hindwing eyespots, the best-fitting model contained treatment, sex, and the interaction between treatment and sex as explanatory variables (Table S2). Although sex had no influence on eyespot size overall (p = 0.483), female individuals for which the humidity was switched from 60% to 85% at the prepupal stage had larger relative hindwing eyespot size compared to those eclosed from the *60%* treatment (Fig. 3B; p = 0.012).

**Figure 3.**
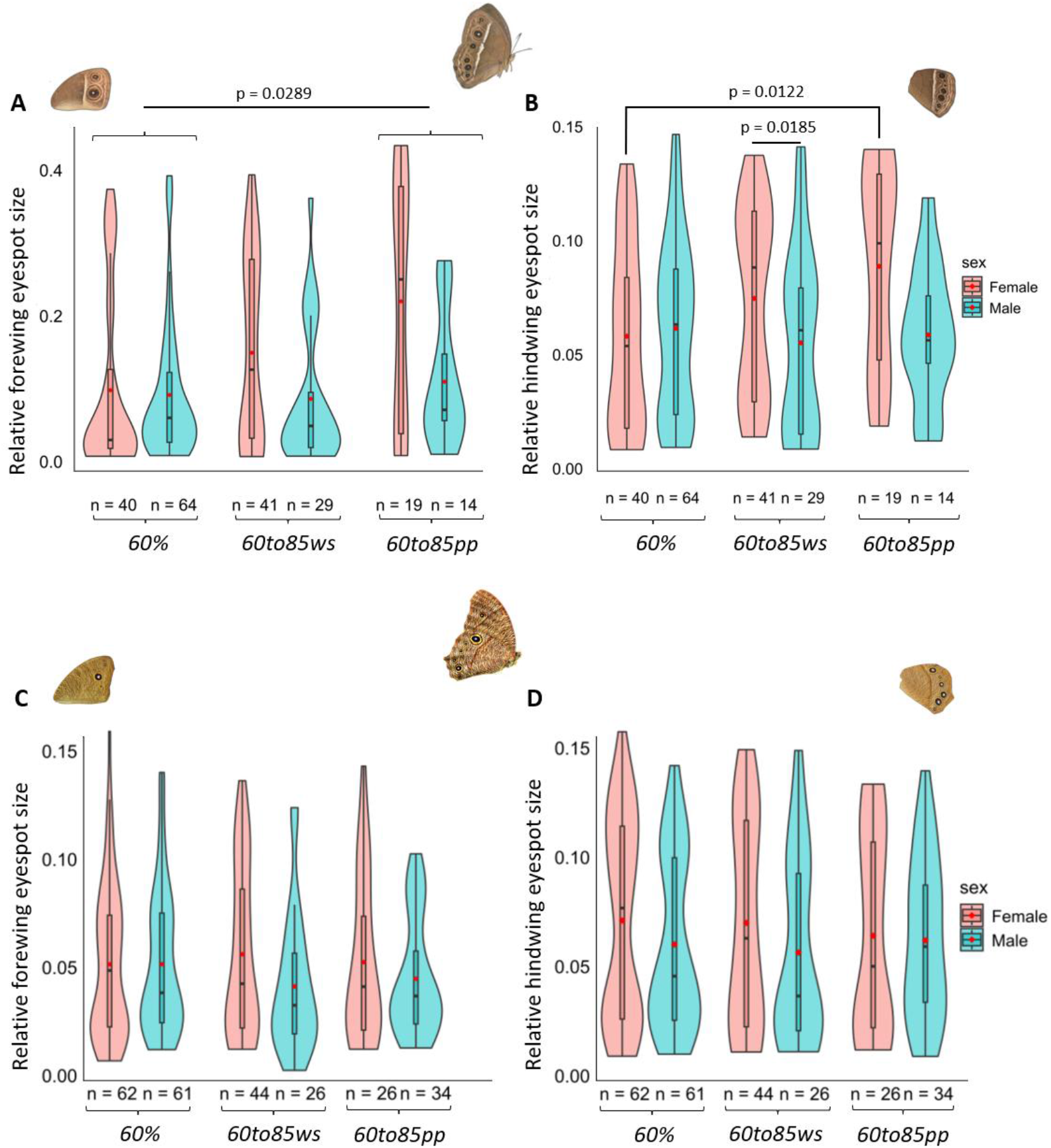
Violin plots showing the effects of switching humidity at multiple life stages from 60% to 85% on the relative eyespot sizes of (A) forewing and (B) hindwing in *My. mineus*, and (C) forewing and (D) hindwing in *Me. leda*. The boxes inside the violin represent the 75^th^ (upper horizontal lines) and 25^th^ (lower horizontal lines) percentiles of the samples and medians (middle horizontal lines). The whisker lines indicate 1.5 times the interquartile range below and above the 25^th^ and 75^th^ percentiles, respectively. The red circles represent the respective averages. Shaded regions surrounding the boxes depict rotated kernel density plots. P-values reported are based on Tukey’s *post hoc* tests. Only significant differences between treatments are shown.

Among the GLMs to test the effect of switching humidity from 85% to 60% on relative forewing eyespot size of *My. mineus*, the best-fitting model had treatment as the sole explanatory variable (Table S3), similar to the opposite switch described above. Adults from the *85%* humidity treatment had larger relative forewing eyespot size compared to adults from the *85to60ws* (Tukey’s *post hoc* test: p < 0.0001) and *85to60pp* (Tukey’s *post hoc* test: p < 0.0001); however, there was no difference between *85to60ws* and *85to60pp* (Fig. 4A; Tukey’s *post hoc* test: p = 0.952). Results were similar for the relative hindwing eyespot size, with treatment being the only significant explanatory variable (Table S4). There was no difference in the relative eyespot size between *80to60ws* and *80to60pp* (Fig. 4B; Tukey’s *post hoc* test: p = 0.974). However, adults from *85%* humidity had significantly larger eyespot sizes compared to those from *80to60ws* and *80to60pp* (Fig. 4B; Tukey’s *post hoc* test: p < 0.0001 for both comparisons). Overall, the results of the two switching directions were similar, except that there was no effect when switching from low to high during the wandering stage, while there was such an effect when switching from high to low humidity. Note that individuals switched during the wandering stage remained at the new humidity during the prepupal stage as well.

**Figure 4.**
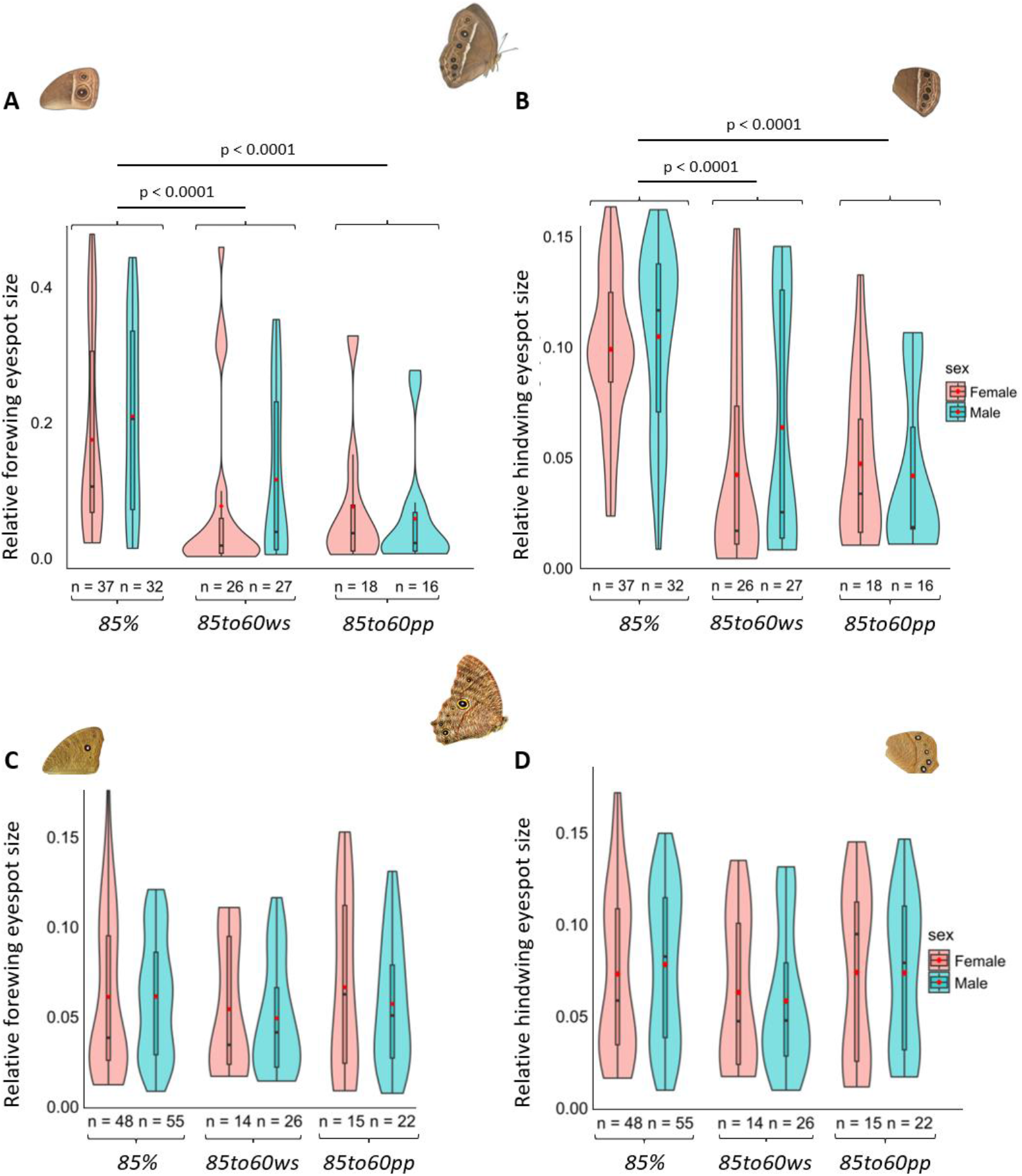
Violin plots showing the effects of switching humidity at multiple life stages from 85% to 60% on the relative eyespot sizes of (A) forewing and (B) hindwing in *My. mineus*, and (C) forewing and (D) hindwing in *Me. leda*. The boxes inside the violin represent the 75^th^ (upper horizontal lines) and 25^th^ (lower horizontal lines) percentiles of the samples and medians (middle horizontal lines). The whisker lines indicate 1.5 times the interquartile range below and above the 25^th^ and 75^th^ percentiles, respectively. The red circles represent the respective averages. Shaded regions surrounding the boxes depict rotated kernel density plots. P-values reported are based on Tukey’s *post hoc* tests. Only significant differences between treatments are shown.

#### ii) Melanitis leda

For both forewing and hindwing eyespot size, the null model was the best-fitting model in both humidity switching experiments from 60% to 85% (Table S5) and 85% to 60% (Table S6). The relative forewing eyespot size showed no significant difference across humidity switching treatments from 60% to 85% (Fig. 3C; p-values based on Tukey’s *post hoc* test: *60%* vs *60to85ws* = 0.770, *60%* vs *60to85pp* = 0.875 and *60to85ws* vs *60to85pp* = 0.988). Similarly, there was no significant difference in the relative hindwing eyespot size across the humidity switching treatments (Fig. 3D; p values based on Tukey’s *post hoc* test: *60%* vs *60to85ws* = 0.907, *60%* vs *60to85pp* = 0.939 and *60to85ws* vs *60to85pp* = 0.998).

For switching treatments from 85% to 60% humidity, there was no significant difference in the relative forewing eyespot size across the humidity switching treatments in pairwise comparisons (Fig. 4C; p values based on Tukey’s *post hoc* test: *85%* vs *85to60ws* = 0.435, *85%* vs *85to60pp* = 0.928 and *85to60ws* vs *85to60pp* = 0.760). Similarly, there was no difference across treatments in the relative hindwing eyespot area (Fig. 4D; p values based on Tukey’s *post hoc* test: *85%* vs *85to60ws* = 0.132, *85%* vs *85to60pp* = 0.928 and *85to60ws* vs *85to60pp* = 0.401).

### Quantification of host plant quality

As a measure of host plant quality, the C:N ratio was quantified for the three host plants used in *60lowhost, 60%* and *85%* humidity treatments. Drought-stressed plants in *60lowhost* had the highest C:N ratio. C:N ratio was higher in *60lowhost* than in both *60%* (Fig. 5; Dunn’s *post hoc* test: p = 0.035) and *85%* humidity (Fig. 5; Dunn’s *post hoc* test: p < 0.0001). C:N was higher in *60%* than in *85%* humidity (Fig. 5; Dunn’s *post hoc* test: p < 0.0001).

**Figure 5.**
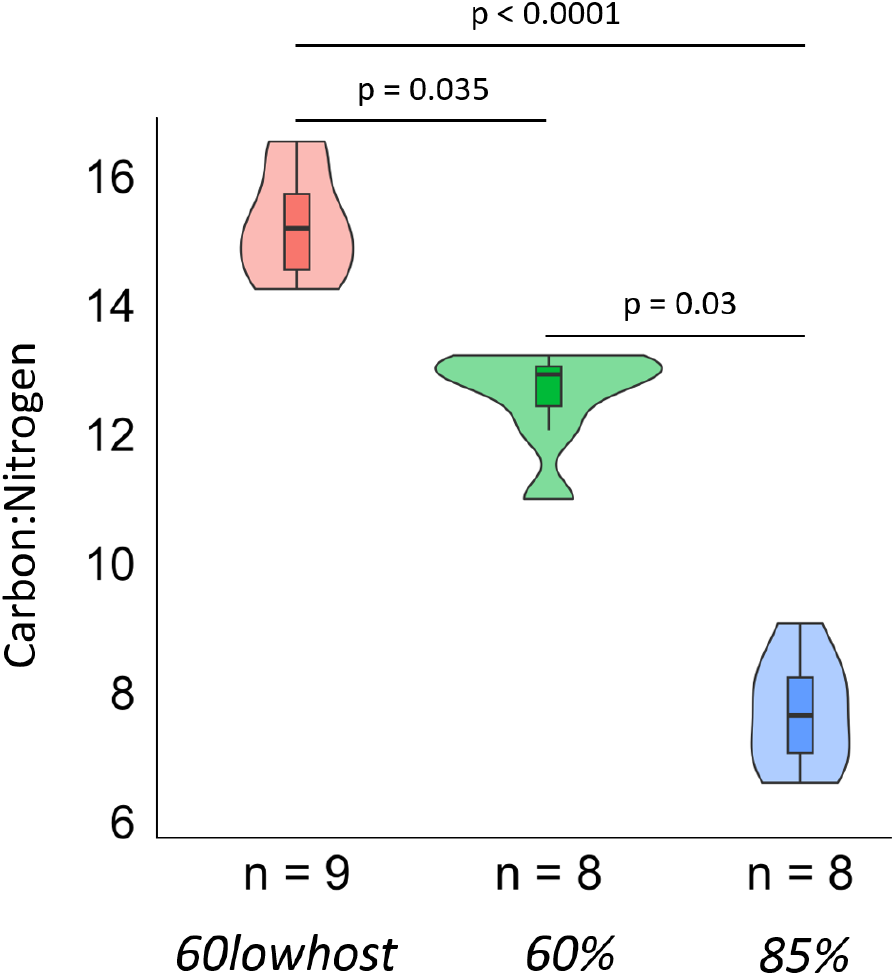
Violin plots showing the Carbon:Nitrogen of the host plants used in *60lowhost* (pink), *60%* (green) and *85%* (blue) humidity treatments to test whether humidity-induced changes in host plant quality affect eyespot size. The boxes inside the violin represent the 75^th^ (upper horizontal lines) and 25^th^ (lower horizontal lines) percentiles of the samples and medians (middle horizontal lines). The whisker lines indicate 1.5 times the interquartile range below and above the 25^th^ and 75^th^ percentiles, respectively. Shaded regions surrounding the boxes depict rotated kernel density plots. P-values reported are based on Dunn’s *post hoc* tests with Holm corrections.

### Comparison 3: Effect of humidity via host plant

#### i) Mycalesis mineus

For *My. mineus*, the best-fitting models to test the indirect effect of humidity via induced changes in the host plant on eyespot sizes for both forewing and hindwing contained treatment as the only explanatory variable. Adults developed at *85%* humidity treatment had significantly larger relative forewing eyespot size compared to adults reared at 60% humidity on plants that had been kept at low humidity (*60lowhost*; Tukey’s *post hoc* test: p < 0.0001) and well-watered plants (*60%:* Tukey’s *post hoc* test: p < 0.0001). There was no difference between the *60lowhost* and *60%* humidity (Fig. 6A; Tukey’s *post hoc* test: p = 0.205). Similarly, hindwing eyespot sizes were larger at *85%* humidity than at *60lowhost* (Tukey’s *post hoc* test: p < 0.0001) and *60%* humidity (Tukey’s *post hoc* test: p < 0.0001), while eyespot sizes did not differ between *60lowhost* and *60%* humidity treatments (Fig. 6B; Tukey’s *post hoc* test: p = 0.260).

**Figure 6.**
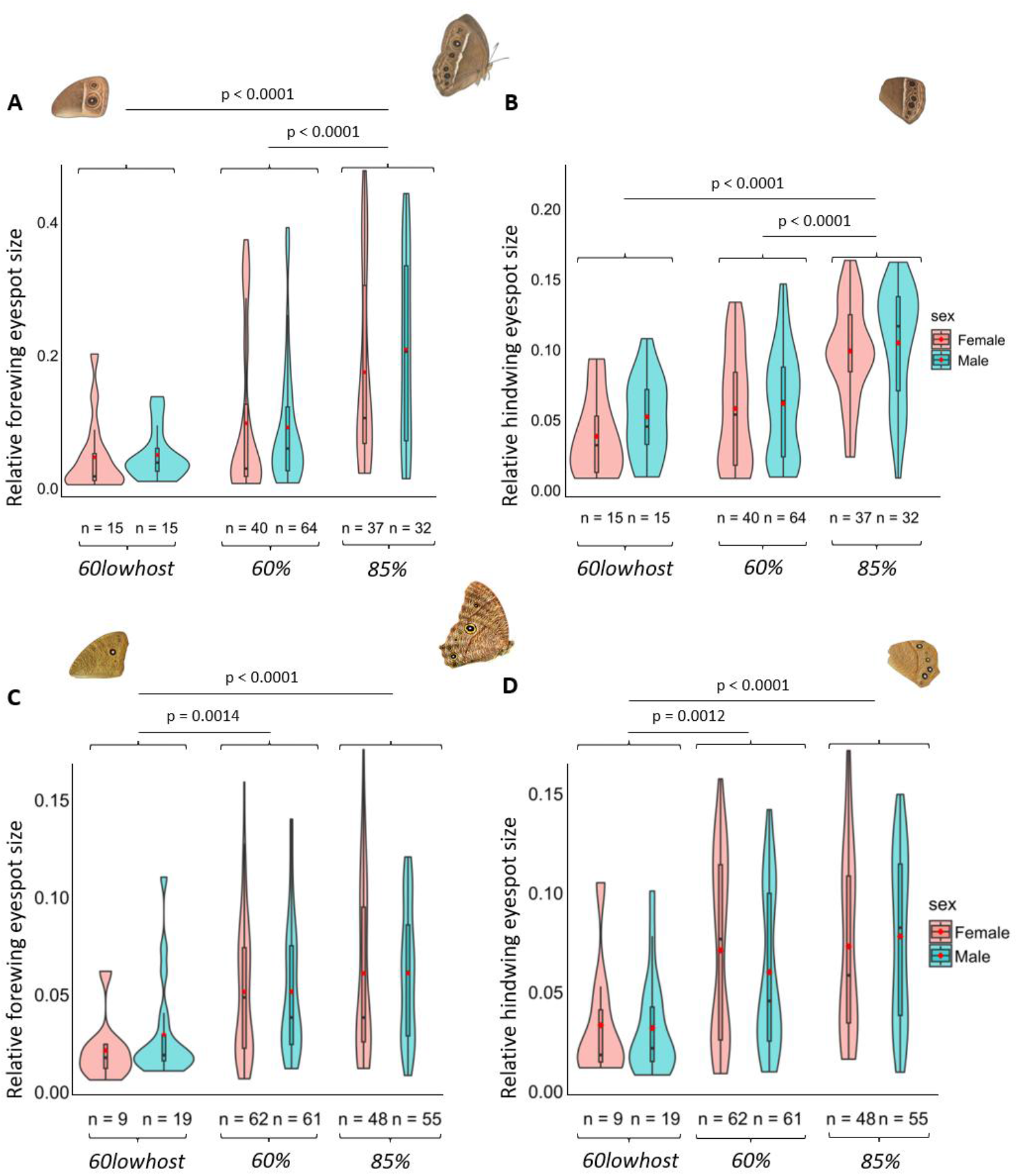
Violin plots showing the effects of host plants grown at different humidity on the relative eyespot sizes of (A) forewing and (B) hindwing in *My. mineus*, and (C) forewing and (D) hindwing in *Me. leda*. The boxes inside the violin represent the 75^th^ (upper horizontal lines) and 25^th^ (lower horizontal lines) percentiles of the samples and medians (middle horizontal lines). The whisker lines indicate 1.5 times the interquartile range below and above the 25^th^ and 75^th^ percentiles, respectively. The red circles represent the respective averages. Shaded regions surrounding the boxes depict rotated kernel density plots. P-values reported are based on Tukey’s *post hoc* tests. Only significant differences between treatments are shown.

#### ii) Melanitis leda

The best-fitting models contained treatment as the sole explanatory variable for both forewing and hindwing in *Me. leda*. Adults from *60lowhost* had significantly smaller relative forewing eyespot sizes than those from *60%* (Tukey’s *post hoc* test: p = 0.001) and *85%* humidity (Tukey’s *post hoc* test: p < 0.0001). Similarly, the relative hindwing eyespot sizes were significantly smaller at *60lowhost* than at *60%* (Tukey’s *post hoc* test: p = 0.001) and *85%* humidity (Fig. 6D; Tukey’s *post hoc* test: p < 0.0001). There was no difference in eyespot size between *60%* and *85%* humidity on both forewing (Fig. 6C; Tukey’s *post hoc* test: p = 0.164) and hindwing (Fig. 6D; Tukey’s *post hoc* test: p = 0.100).

## Discussion

We investigated how humidity may be used as a cue for seasonal polyphenism in sympatric populations of two tropical satyrine butterfly species from distinct lineages that share climate, habitat and host plants. In *Mycalesis mineus*, we found that humidity directly influenced eyespot size on both fore- and hindwings (Fig. 2A, B) and that the humidity-sensitive developmental period occurs during the wandering and prepupal stages (Fig. 3A, B and Fig. 4A, B). In contrast, in *Melanitis leda*, humidity did not modulate eyespot size directly (Fig. 2C, D), but this species responded strongly to host plant quality (Fig. 6C, D). The host plant mediated responses to humidity fitted with host plant quality differences (Fig. 5), with larvae that fed on lower quality plants (high Carbon:Nitrogen) tending to have smaller eyespots as adults.

### Direct and indirect effects of humidity

Several studies have pointed out that cues other than temperature are likely needed to explain seasonal polyphenism of tropical satyrines in regions where temperature is a poor predictor of precipitation (11,34,35). In this study, we show that humidity, which increases with the onset of the wet season and decreases towards the end of the wet season, can be used as a cue for eyespot size plasticity in two distantly related lineages, suggesting that using humidity as a cue for seasonal polyphenism may be widespread. Notably, the mechanisms through which the two species used humidity as a cue differ: *My. mineus* uses humidity sensed during the larval stage (Fig. 2A, B), while *Me. leda* only responded to differences between well-watered and drought-stressed plants (Fig. 6C, D). The Carbon-Nitrogen ratio analyses confirm that the well-watered plants were of higher quality (Fig. 5). Previous work showed that host plant species affect eyespot size in both of our study species and *Ypthima huebneri* (4,23), and *B. anynana* (same sub-tribe as *My. mineus*) (45). Moreover, drought treatment (48) and larval food limitation (68) can induce smaller eyespots in *B. anynana*. Therefore, using host plant quality as a cue for seasonal polyphenism may be widespread. However, more studies are needed to establish how general this pattern is, and what mechanisms underlie it.

Using humidity as a direct cue (during non-feeding immature stages) for eyespot size appears to be less common (one out of the two investigated lineages) than indirect effects via the host plant. Given that humidity is often closely correlated with precipitation, we may expect humidity to be a widely used cue, but most comparative studies have focused on temperature. Notably, the critical period for humidity appears to coincide with the critical period for temperature in *B. anynana*; the late larval and early pupal stages. To our knowledge, this is the first study that maps a non-temperature cue to a specific developmental period in butterflies. This period is probably most predictive of the adult environment, and just before wing traits are developing. Further studies are needed to establish how widespread direct humidity cue use is across butterfly lineages, and if the critical period is always during these non-feeding stages.

Since both the larvae and the host plant on which larvae feed are under the same conditions inside the growth chamber during an experiment, it is virtually impossible to completely segregate the direct effects of the environment on the larvae from those via host plants. Especially the indirect effect via host plant quality may be complex, as the response of the plants may depend on the environment they were grown in before being placed in the growth chamber. Thus, a plant acclimated to low humidity may suffer when placed at high humidity, and *vice versa*. This is not unique to humidity: plants will also respond to temperature. Thus, even if host plant quality appears to be constant, the environment under which the plant was grown will affect how it responds to being placed under different environmental conditions. We tried to avoid this effect by conducting trials across all seven treatments simultaneously, such that they all used the same batches of plants. Furthermore, to limit the effects of humidity on host plants, we replaced them every two days, which is quite frequent compared to other experiments in which plants are replaced ‘when needed’ (e.g., 4,26); nevertheless, an effect of humidity on host plant quality was evident (Fig. 5). Direct cue-use appears to be concentrated after the larval feeding phase (wandering and prepupal stages; our results for humidity, and Kooi & Brakefield (29) and Monteiro et al. (30) for temperature), making it relatively easy to distinguish these direct cues from indirect effects via the plant. However, this may not be the case for all species. Apart from carefully considering these indirect effects in experimental design, future studies would also benefit from measuring the quality of plants used in experiments, as well as including time series of plant quality in field studies (35). Finally, it appears from our study that the Carbon-Nitrogen ratio is a good proxy for plant quality in this system, but other plant traits may also be important.

## Conclusion

Although there is substantial evidence that temperature influences eyespot size in butterflies with dry-wet season polyphenism in tropical satyrines, temperature is not always a reliable cue in nature, so other cues must be used. This study shows that butterflies can use humidity, which is seasonally coupled with rainfall, to modulate eyespot size plasticity. We demonstrate that in *My. mineus* humidity mainly directly affects eyespot size, with the critical period of sensitivity occurring during the late larval and prepupal stages when feeding on host plants is completed. In contrast, in *Me. leda*, humidity affects the eyespot size indirectly through humidity-mediated changes in host plant quality. This shows that different lineages can use humidity in distinct ways to achieve seasonal polyphenism, despite evolving under similar ecological pressures. By using multiple cues or cues that are more consistently linked to seasonal rainfall (like humidity and host plant quality), species that use seasonal polyphenism may be better prepared to cope with or adapt to climate change, than when using a single cue such as temperature.

## Acknowledgement

We thank Hazekaiah Graham Laloo for his contribution during pilot experiments, and Nikhil Kollins for his help in rearing the experimental population and collecting part of the data of *Myclaesis mineus*. We thank members of Vanasiri Evolutionary Ecology Lab (www.vanasiri.in) – Akhil Sadiq, Anaswara K. Sreedharan, Aarini Ghosh, Bhanu B. Sharma, Indukala Prasannakumar, Kushankur Bhattacharyya, Sharafudheen Thangal, Shreya Mishra and – for their comments on the manuscript and their help in the collection and rearing of the butterflies.

