## Supplementary material for "Contrasting mechanisms for using humidity as cue for seasonal polyphenism in two tropical butterflies": Table

**Table S1.** Regression coefficients of the best-fitting model from *glmmTMB* analyses to test the effect of switching humidity from 60% to 85% on the relative forewing eyespot size in *Mycalesis mineus*. SE and $R^2$ represent the standard error of the factors and the Ferrari's R-square of the model, respectively.

| Factor | Estimate | SE | t value | p value | AIC | $R^2$ |
|---|---|---|---|---|---|---|
| Intercept | -2.159 | 0.100 | -21.624 | < 0.0001 | -478.7 | 0.036 |
| treatment60to85pp | 0.436 | 0.171 | 2.552 | 0.011 | | |
| treatment60to85ws | 0.157 | 0.137 | 1.142 | 0.2535 | | |

**Table S2.** Regression coefficients of the best-fitting model from *glmmTMB* analyses to test the effect of switching humidity from 60% to 85% on the relative hindwing eyespot size in *My. mineus*. SE and $R^2$ represent the standard error of the factors and the Ferrari's R-square of the model, respectively.

| Factor | Estimate | SE | t value | p value | AIC | $R^2$ |
|---|---|---|---|---|---|---|
| Intercept | -2.819 | 0.109 | -25.934 | < 0.0001 | -771.0 | 0.058 |
| Treatment: 60to85pp | 0.482 | 0.170 | 2.849 | 0.004 | | |
| Treatment: 60to85ws | 0.305 | 0.142 | 2.140 | 0.032 | | |
| Sex Male | 0.094 | 0.134 | 0.701 | 0.483 | | |
| Treatment: 60to85pp: sex Male | -0.460 | 0.255 | -1.804 | 0.071 | | |
| Treatment: 60to85ws: sex Male | -0.468 | 0.208 | -2.254 | 0.024 | | |

**Table S3.** Regression coefficients of the best-fitting model from *glmmTMB* analyses to test the effect of switching humidity from 85% to 60% on the relative forewing eyespot size in *My. mineus*. SE and $R^2$ represent the standard error of the factors and the Ferrari's R-square of the model, respectively.

| Factor | Estimate | SE | t value | p value | AIC | $R^2$ |
|---|---|---|---|---|---|---|
| Intercept | -1.452 | 0.104 | -13.995 | < 0.0001 | -362.6 | 0.24 |
| Treatment: 60to85pp | -0.959 | 0.192 | -4.993 | < 0.0001 | | |
| Treatment: 60to85ws | -0.870 | 0.165 | -5.442 | < 0.0001 | | |

**Table S4**. Regression coefficients of the best-fitting model from *glmmTMB* analyses to test the effect of switching humidity from 85% to 60% on the relative hindwing eyespot size in *My. mineus*. SE and $R^2$ represent the standard error of the factors and the Ferrari's R-square of the model, respectively.

| **Factor** | **Estimate** | **SE** | **t value** | **p value** | **AIC** | **$R^2$** |
|---|---|---|---|---|---|---|
| Intercept | -2.138 | 0.068 | -31.296 | < 0.0001 | -567.3 | 0.30 |
| Treatment: 85to60pp | -0.879 | 0.143 | -6.166 | < 0.0001 | | |
| Treatment: 85to60ws | -0.914 | 0.122 | -7.465 | < 0.0001 | | |

**Table S5.** Model selection from *glmmTMB* analyses to test the effect of switching humidity from 60% to 85% on the relative forewing (FW) and hindwing (HW) eyespot size in *Melanitis leda* based on likelihood ratio tests using Chi-squared statistics (Chi sq.) and AICc. LogLik represents log likelihood. Best-fitting model is highlighted in bold.

| **Model number** | **Model** | **Comparison** | **LogLik** | **Chi sq.** | **AICc** | **p value** |
|---|---|---|---|---|---|---|
| FOREWING (FW) EYESPOT SIZE | | | | | | |
| Fw0 | FW ~ $(\text{treatment + sex})^2$ | Fw0, Fw1 | 543.68 | 2.824 | -1072.905 | 0.244 |
| Fw1 | FW ~ treatment + sex | Fw1, Fw2 | 542.27 | 0.852 | -1074.295 | 0.356 |
| Fw1 | FW ~ treatment + sex | Fw1, Fw3 | 542.27 | 0.575 | -1074.295 | 0.750 |
| Fw2 | FW ~ treatment | Fw2, Fw4 | 541.84 | 0.552 | -1075.525 | 0.759 |
| Fw3 | FW ~ sex | Fw3, Fw4 | 541.98 | 0.830 | -1077.867 | 0.362 |
| **Fw4** | **FW ~ 1 (Null model)** | | **541.57** | | **-1079.086** | |
| HINDWING (HW) EYESPOT SIZE | | | | | | |
| Hw0 | HW ~ $(\text{treatment + sex})^2$ | Hw0, Hw1 | 470.53 | 0.812 | -926.604 | 0.666 |
| Hw1 | HW ~ treatment + sex | Hw1, Hw2 | 470.12 | 1.201 | -930.006 | 0.273 |
| Hw1 | HW ~ treatment + sex | Hw1, Hw3 | 470.12 | 0.297 | -930.006 | 0.862 |
| Hw2 | HW ~ treatment | Hw2, Hw4 | 469.52 | 0.221 | -930.887 | 0.895 |
| Hw3 | HW ~ sex | Hw3, Hw4 | 469.98 | 1.125 | -933.855 | 0.289 |
| **Hw4** | **HW ~ 1 (Null model)** | | **469.41** | | **-934.779** | |

**Table S6.** Model selection from *glmmTMB* analyses to test the effect of switching humidity from 85% to 60% on the relative forewing (FW) and hindwing (HW) eyespot size in *Me. leda* based on likelihood ratio tests using Chi-squared statistics (Chi sq.) and AICc. LogLik represents log likelihood.

| Model number | Model | Comparison | LogLik | Chi sq. | AICc | p value |
|---|---|---|---|---|---|---|
| FOREWING (FW) EYESPOT SIZE | | | | | | |
| Fw0 | FW ~ (treatment + sex)$^2$ | Fw0, Fw1 | 359.77 | 0.456 | -704.885 | 0.796 |
| Fw1 | FW ~ treatment + sex | Fw1, Fw2 | 359.54 | 0.115 | -708.736 | 0.734 |
| Fw1 | FW ~ treatment + sex | Fw1, Fw3 | 359.54 | 1.645 | -708.736 | 0.440 |
| Fw2 | FW ~ treatment | Fw2, Fw4 | 359.48 | 1.563 | -710.737 | 0.458 |
| Fw3 | FW ~ sex | Fw3, Fw4 | 358.72 | 0.035 | -711.301 | 0.852 |
| **Fw4** | **FW ~ 1 (Null model)** | | **358.70** | | **-713.335** | |
| HINDWING (HW) EYESPOT SIZE | | | | | | |
| Hw0 | HW ~ (treatment + sex)$^2$ | Hw0, Hw1 | 328.50 | 0.306 | -642.339 | 0.858 |
| Hw1 | HW ~ treatment + sex | Hw1, Hw2 | 328.34 | 0.341 | -646.339 | 0.560 |
| Hw1 | HW ~ treatment + sex | Hw1, Hw3 | 328.34 | 4.120 | -646.339 | 0.128 |
| Hw2 | HW ~ treatment | Hw2, Hw4 | 328.17 | 3.899 | -648.115 | 0.142 |
| Hw3 | HW ~ sex | Hw3, Hw4 | 326.28 | 0.12 | -646.428 | 0.729 |
| **Hw4** | **HW ~ 1 (Null model)** | | **326.22** | | **-648.377** | |
